# Reconstructing mutational lineages in breast cancer by multi-patient-targeted single cell DNA sequencing

**DOI:** 10.1101/2021.11.16.468877

**Authors:** Jake Leighton, Min Hu, Emi Sei, Funda Meric-Bernstam, Nicholas E. Navin

## Abstract

Single cell DNA sequencing (scDNA-seq) methods are powerful tools for profiling mutations in cancer cells, however most genomic regions characterized in single cells are non-informative. To overcome this issue, we developed a Multi-Patient-Targeted (MPT) scDNA-seq sequencing method. MPT involves first performing bulk exome sequencing across a cohort of cancer patients to identify somatic mutations, which are then pooled together to develop a single custom targeted panel for high-throughput scDNA-seq using a microfluidics platform. We applied MPT to profile 330 mutations across 23,500 cells from 5 TNBC patients, which showed that 3 tumors were monoclonal and 2 tumors were polyclonal. From this data, we reconstructed mutational lineages and identified early mutational and copy number events, including early *TP53* mutations that occurred in all five patients. Collectively, our data suggests that MPT can overcome technical obstacles for studying tumor evolution using scDNA-seq by profiling information-rich mutation sites.

## Introduction

Triple negative breast cancer (TNBCs) is an aggressive subtype that is characterized by a lack of estrogen receptor (ER), progesterone receptor (PR) and Her2 amplification. TNBC patients frequently develop resistance to chemotherapy (~50%) and progress to metastatic disease, often leading to poor survival (Foulkes et al. 2010; Liedtke et al. 2008; Kim et al. 2018). In contrast to ER-positive breast cancer, TNBC patients display extensive copy number alterations (CNAs) and driver mutations in *TP53* in 83% of patients (TCGA, 2012). Furthermore, TNBC patients have been shown by single cell DNA sequencing (scDNA-seq), multi-region sequencing and whole-genome sequencing methods to display extensive intratumor heterogeneity (ITH) (Gao et al. 2016; Yates et al. 2015; Shah et al. 2012). This ITH may explain why TNBC patients frequently develop resistance to chemotherapy and often progress to metastatic disease (Nedeljkovoc et al. 2019; Marra et al. 2020). Previous studies have shown that TNBCs evolve through Punctuated Copy Number Evolution (PCNE) in which large numbers of CNA events are acquired in short bursts of evolution at the earliest stages of progression (Gao et al. 2016; Kim et al. 2018; Casasent et al. 2018; Minussi et al. 2021). However, the dynamics and timing of mutations during TNBC progression represent a fundamental gap in knowledge. Additionally, improved knowledge of ITH is important for the clinical diagnosis and treatment of TNBC patients, particularly when considering targeted therapy. One approach for resolving intratumor heterogeneity that is gaining broader use in both research and clinical studies is scDNA-seq (Lim et al. 2020).

The field of single cell genomics has shown remarkable progress in the development of both transcriptomic and genomic profiling methods over the last 10 years (Navin 2014). Initial single cell DNA sequencing methods (scDNA-seq) for profiling the genomes and exomes of single cells were low-throughput and had extensive technical errors (Navin et al. 2011; Zong et al. 2021; Wang et al. 2014). These technologies have been improved through developments in whole genome amplification (WGA) chemistries (Telenius et al. 1992; Dean et al. 2002; Zong et al. 2012; Kamberov et al. 2012) and the implementation of nanowell and microfluidic platforms (Gierahn et al. 2017; Prakaden et al. 2017; Macosko et al. 2015, Nagasawa et al. 2021). Specifically, single cell genomic copy number profiling technologies have undergone substantial improvements with the use of tagmentation chemistries and microfluidic or nanowell platforms (Minussi et al. 2021; 10X Genomics CNV, Laks et al. 2019). Similarly, scDNA-seq platforms for mutational profiling have been developed using microfluidic platforms (Mission Bio, Tapestri) (Lan et al. 2017; Pelligrino et al. 2018; Morita et al. 2020; Miles et al. 2020; Wang et al. 2021), which have enabled the profiling of tens of thousands of cells in parallel. However, in contrast to genomic copy number profiling, which involves unbiased whole genome sequencing (WGS) at sparse depth, mutational profiling requires high-coverage data. Consequently, it is necessary to target genomic regions for sequencing (eg. exome, cancer gene panels) in order to make the assays economically feasible. This presents a major challenge, since the informative mutation sites are not usually known *a priori*, and thus most of the genomic regions profiled in single cells contain only reference DNA sequences or SNPs.

To address this challenge, we developed a Multi-Patient-Targeted (MPT) scDNA-seq sequencing approach that combines bulk DNA sequencing, single-cell droplet-based microfluidics, and a custom targeted panel. We applied MPT to 5 invasive TNBC tumor samples, which resolved the clonal substructure of the tumors and enabled the reconstruction of clonal lineages during tumor progression. Our data show that *TP53* mutation and other early mutational events in TNBC progression are accompanied by CNA events, leading to the expansion of the primary tumor mass. Additionally, our data show that most mutations are early events in the progression of TNBC tumors, and that subclones undergo stable clonal expansions with only limited intermediate mutations that are acquired during tumor growth.

## Results

### Multi-Patient-Targeted Panel (MPT) Sequencing

A major challenge with targeted scDNA-seq of mutations in tumors, is that generally only a few somatic mutations (eg. 1-10) can be profiled from each patient using an unbiased panel that covers a small region (eg. 5mb) of the human genome, while most genomic regions sequenced contain only the reference genome bases and SNPs. This problem results in high costs for scDNA-seq of regions with limited useful mutational information and prohibits large numbers of cell to be profiled in parallel. To address this problem, we developed a method called Multi-Patient-Targeted (MPT) sequencing (Fig. 1). In this approach, we first profile a set of patients (eg. 5-20) with bulk deep-exome sequencing methods (approximately ~200X tumor, 60X normal) to identify somatic mutations and then pool all mutations together from the patient cohort (eg. ~1000 mutations) to develop a custom targeted panel for scDNA-seq that targets all of the mutation sites across all of the patients (Fig. 1A). The custom panel is synthesized commercially (V1 custom targeted panel, Mission Bio) for the purpose of downstream scDNA-seq using a microfluidics platform (Tapestri, Mission Bio) (Fig. 1B). We then perform scDNA-seq (Mission Bio) using the same MPT panel on each of the tumors that were previously analyzed by bulk exome sequencing to profile ~5000 cells for each tumor (Fig. 1C). In the scDNA-seq experiments that are performed for each tumor, the mutations are profiled for each patient (eg. 200 mutations), as well as the reference sites for the mutations in the other patients (which are not utilized in the downstream analysis). The major advantage of the MPT approach is that it greatly mitigates the sequencing space for non-informative genome information that would normally be profiled using an unbiased panel. By focusing the scDNA-seq on the mutation sites of interest, the sequencing costs are greatly decreased, enabling very high-throughput analysis of thousands of single cells in parallel at low costs for each patient.

**Figure 1.**
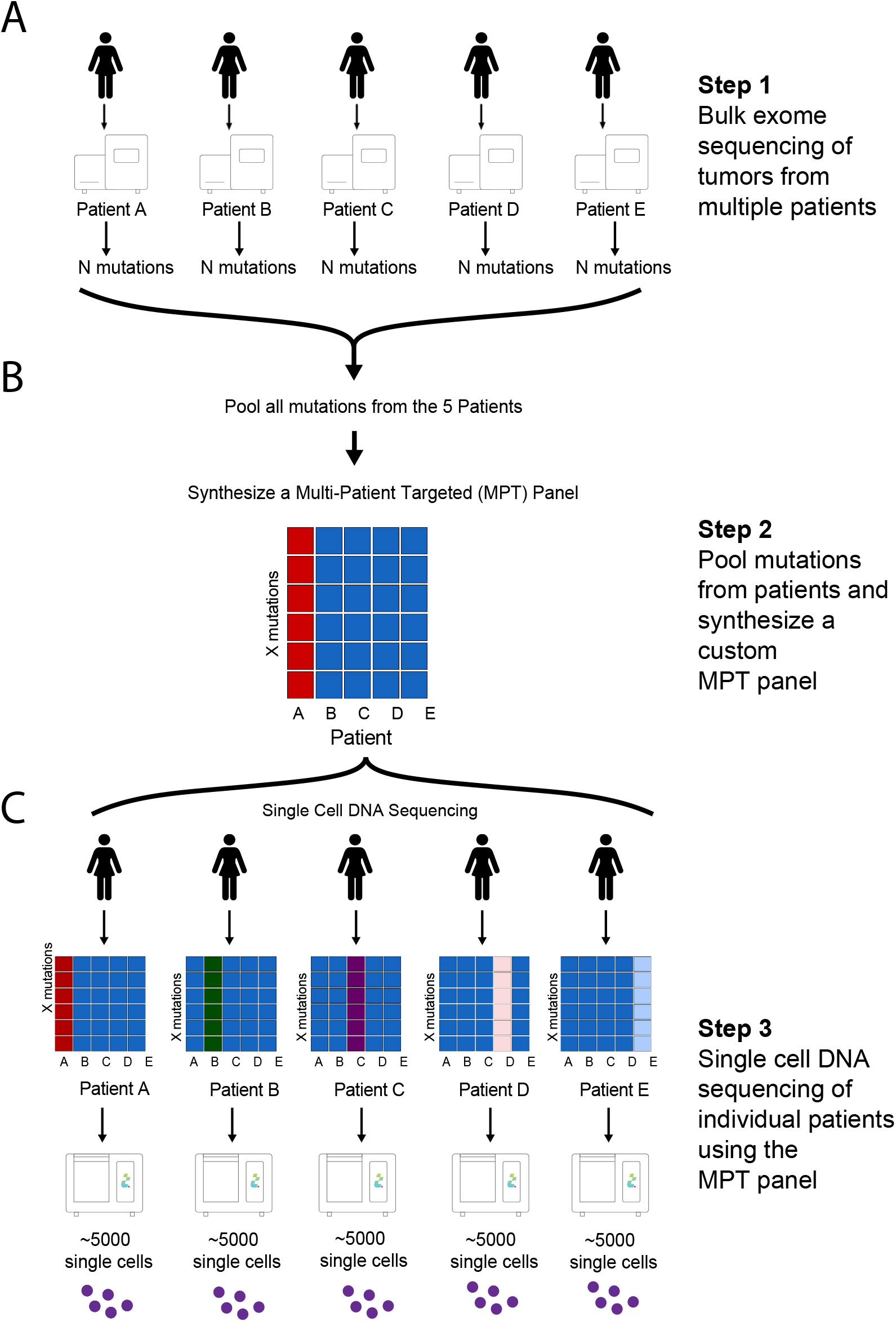
Overview of Multi-Patient-Targeted Single Cell Sequencing Workflow. (A) Bulk exome sequencing is performed on a cohort of human tumors (B) Mutations from all patients are pooled together and used to synthesize a MPT custom targeted panel (C) The MPT panel is used to perform scDNA-seq on each patient individually to profile thousands of cells in parallel using a microfluidics platform.

### Bulk Exome Sequencing of Triple-Negative Breast Cancers

We selected frozen invasive tumor samples from 5 untreated TNBC patients that were negative for the expression of estrogen, progesterone and Her2 receptors for bulk exome sequencing followed by MPT profiling using a microfluidics system (Tapestri, Mission Bio) (Supplementary Table 1). To enrich the tumor cell populations, we first generated single nuclei suspensions and stained the nuclei with DAPI to perform fluorescence-activated cell sorting (FACS) (Fig. 2A). Using FACS, we gated distributions of cells from 2.65-3.7(N) ploidy to enrich aneuploid tumor cell fractions. Using the aneuploid cells, we generated sequencing libraries and performed both whole-genome sequencing at sparse depth to estimate genomic copy number and exome capture (Roche, Nimblegen V2) to identify point mutations. The genomic copy number profiles showed extensive aneuploidy and copy number aberrations (CNAs) in all of the tumors, including amplifications of oncogenes such as *MYC*, and deletions of tumor suppressors such as *TP53* (Fig. 2B). Exome sequencing of the aneuploid-sorted nuclei and matched normal tissues identified a range of somatic mutations (mean = 66) in the tumors, in which TN4 and TN5 had highly elevated mutation burden compared to TN1-TN3, suggesting that they are hypermutators. Analysis of the mutational signatures showed that several common breast cancer signatures were prevalent, including Signature 1a/b, Signature 3, and Signature 13 (Fig. 2C). Signature 1a/1b is aging (deamination of methylcytosines), signature 3 is homologous recombination repair deficiency and Signature 13 is APOBEC cytidine deaminases. Finally, we investigated the distribution of somatic mutation frequencies which ranged from ~0.1% to ~99% for each patient, and showed that *TP53* was one of the highest frequency mutations identified in all 5 patients (Fig. 2D). We combined all of the somatic mutations identified across the 5 TNBC patients to design a custom MPT panel (Mission Bio) for scDNA-seq analysis.

**Figure 2.**
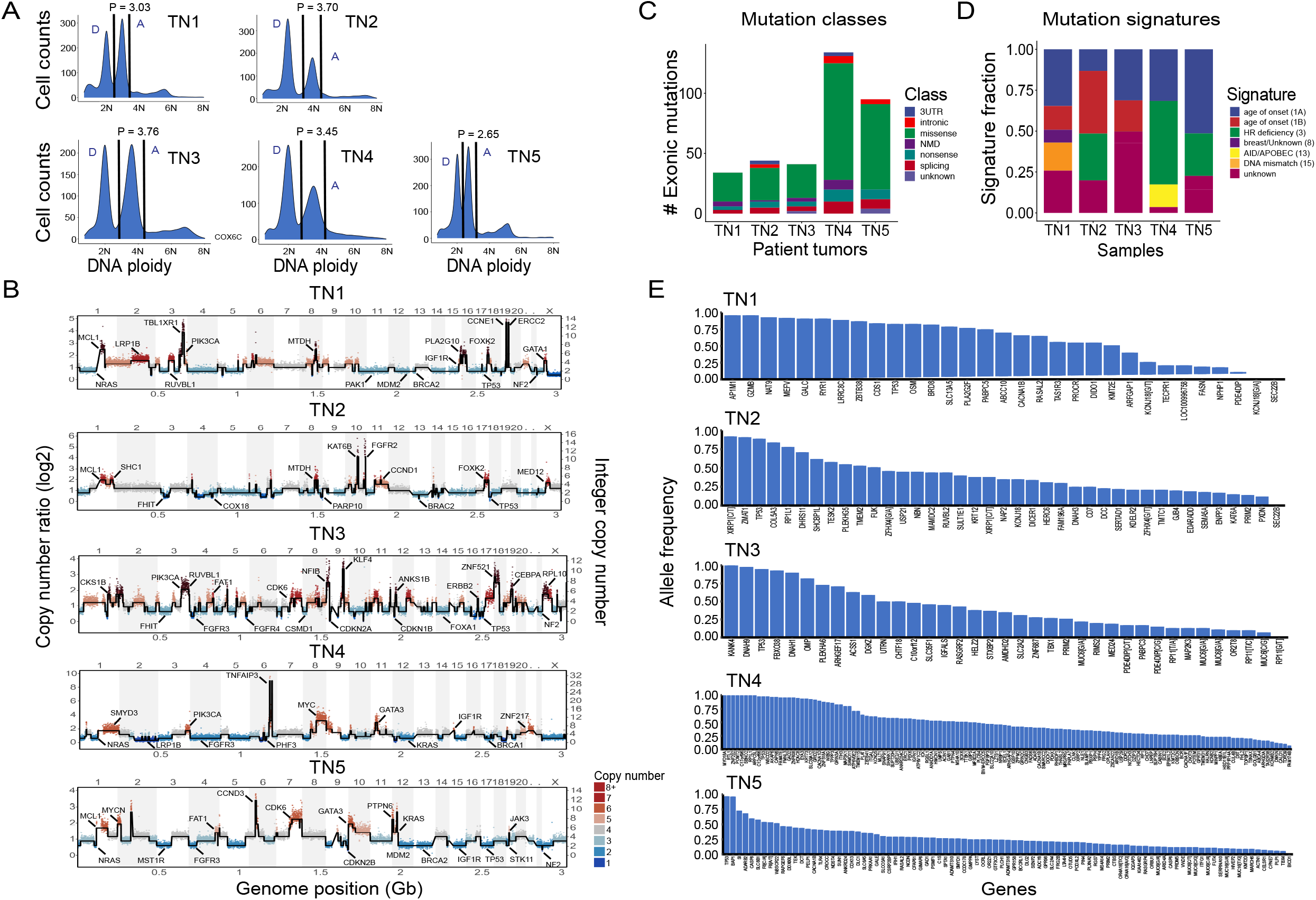
Bulk Exome Sequencing and Mutational Analysis of TNBC Tumors. (A) Frozen tumor tissues were dissociated into nuclear suspensions and stained with DAPI for FACS to isolate aneuploid single cells by differences in DNA ploidy, where P indicates the mean tumor ploidy and D and A indicate diploid or aneuploid distributions (B) Pseudo-bulk copy number ratio plots generated from single cell WGS data from each tumor sample with integer copy number states calculated (C) Classes of exonic mutations for each tumor identified by Annovar (D) Mutational signatures calculated from the exonic mutations (E) Allelic frequencies for all of the somatic mutations identified in each TNBC tumor that were used to design the MPT panel.

### Mutational Substructure of TNBC Tumors

To resolve the clonal substructure of each TNBC tumor, we applied the MPT panel to perform scDNA-seq of a total of 23,526 cells (range 4,002 - 5,941) of the 5 tumors using the microdroplet (Mission Bio) scDNA-seq platform (Supplementary Table 1). The resulting data showed that 330 targeted sites had 164x coverage depth across the 5 tumors. The somatic mutations were detected in each single cell by filtering the matched germline bulk exome sequencing data (Methods).

In the polyclonal tumor TN4, clustering of 4,141 single cells across 69 genes identified 4 major clusters in high-dimensional space using UMAP that corresponded to three tumor subclones (c1-c3) and one population of diploid cells (c4) (Fig. 3A). A clustered heatmap of the mutations further revealed the somatic mutations in each subclone, including shared mutations that were present in all three tumor clusters (Fig. 3B). This analysis identified 15 homozygous mutations that were shared among all three subclones, including *TP53* which is the most commonly mutated gene in TNBC (TCGA 2012). The copy number of each gene on the targeted panel was estimated from the read depth, which identified chromosomal gains and losses (Methods). These data showed that *GRIN3A*, *DMKN* and *IGSF10* were amplified, while *NOTCH3* and 15 other genes with homozygous mutations showed copy number losses. Computational prediction of the functional impact scores using CADD (Kircher et al. 2014) identified *KALRN*, *NOTCH3* and *PTPN4* as having the highest functional impact scores, among other genes (Fig. 3C).

**Figure 3.**
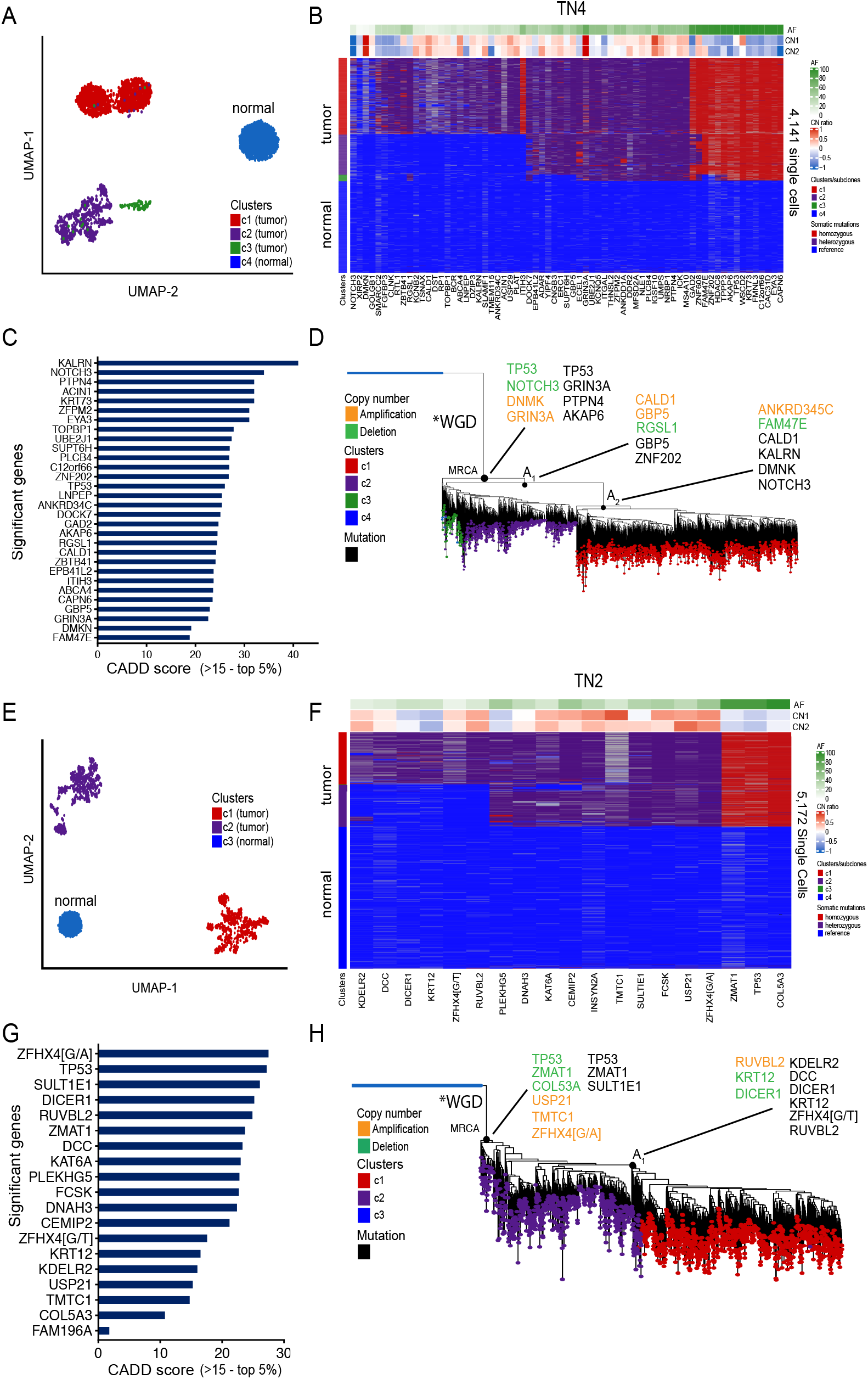
Clonal Substructure and Phylogenetic Reconstruction of Two Polyclonal Tumors. (A, E) High-dimensional clustering of somatic mutations using UMAP of 4,141 and 5,172 (TN2,TN4) single cells to identify major clusters of subclones (B, F) Heatmaps of hierarchical clustered mutations in TN2 and TN4 showing the clonal substructure and heterogeneity within the tumor subpopulations, with cluster-level copy number estimations shown in the header tracks above (C, G) CADD scores are used to rank functional impact of mutations’ deleteriousness in TN2 and TN4 (CADD > 15, top 5%) (D, H) Phylogenetic reconstruction of the mutation lineages in TN2 and TN4, chronology, and timing of copy number aberrations relative to the MRCA using a neighbor-joining tree. The common ancestors (A1, A2) are annotated on the tree, as well as the whole-genome doubling (WGD) events.

To infer the chronology and order of single nucleotide variants (SNVs) and CNAs during tumor evolution, a neighbor-joining (NJ) tree was constructed from the mutation matrix, after which the mutations and CNA events were annotated (Methods). The NJ tree showed three major branching points: the Most Recent Common Ancestor (MRCA), the first subclonal ancestor (A1) and the second subclonal ancestor (A2), which resulted in the divergence of the three tumor subclones (c1-c3). Truncal mutations that occurred in all tumor subclones included chromosome losses in *TP53* and *NOTCH3* and gains in *DNMK* and *GRIN3A*, while early mutations included *TP53*, *GRIN3A*, *PTPN4* and *AKAP6*. The c3 subclone was closest to the MRCA suggesting that it was one of the earliest subclones that diverged, consistent with the clustering results which showed that it harbored the lowest number of mutations (N=33). The c2 clone diverged after the A1 ancestor by acquiring an additional set of mutations (N=8). Finally, the c3 clone diverged from the A2 ancestor via the acquisition of a large set of somatic mutations (N=28) that included a mutation in *NOTCH3* (Fig. 3D). Interestingly, a lack of gradual intermediate mutations was identified between the 3 major subclones in this analysis, and most mutations were very clonal within the three tumor subpopulations, which shared a common evolutionary lineage.

In the polyclonal tumor TN2, a total of 5,172 single cells were sequenced and 19 somatic mutations were identified. Clustering of this data in high-dimensional space identified 3 major clusters, including two major tumor subclones (C1-C2) and one diploid cell cluster (C3) (Fig. 3E). Hierarchical clustering of the somatic mutations identified 13 mutations that were shared between the two tumor clusters including 3 homozygous mutations, of which one was a *TP53* mutation (Fig 3F.). The clustered heatmap also showed 6 mutations that were exclusive to the C2 subclone, suggesting that this subpopulation continued to evolve additional mutations after a shared common ancestor. Applying targeted copy number inference from the read depth data, we identified amplifications in *ZFHX4*[G/A], *USP21* and *TMTC1*, while *TP53*, *ZMAT1* and *COL5A3* showed copy number losses and had homozygous mutations (Fig. 3F). CADD predictions of the functional impact scores showed that *ZFHX4*[G/A], *TP53*, *SULT1E1* and *DICER1* had the highest scores in TN2 (Fig. 3G). A NJ tree was constructed and showed two major branching points in evolution: the Most Recent Common Ancestor (MRCA) and the first subclonal ancestor (A1). Early truncal events that occurred prior to the MRCA included chromosome losses in *TP53*, *ZMAT1* and *COL5A3*, while gains included *USP21*, *TMTC1*, *ZFHX4*[G/A] and mutations included *TP53*, *ZMAT1* and *SULT1E1* (Fig. 3H). The c1 clone diverged from c2 by acquiring 6 additional mutations including *DCC*, *DICER1* and *RUVBL2* with high functional impact scores. Consistent with TN1, the mutational substructure of TN2 showed a lack of gradual mutations detected between the two subclones and the diploid cells.

Clustering of TN1, TN3 and TN5 revealed mono-clonal tumors that were comprised of a single population of tumor cells and a single population of diploid cells (Figure 4). In TN1, 16 mutations were detected across 4,270 single cells. In TN3, the data identified 11 mutations across 5,941 cells while in TN5 47 mutations were detected across 4002 cells (Fig. 4A-B). In TN1, the mutations with the highest impact scores included homozygous mutations in *TP53*, *CDS1*, *RASAL2* and *ARFGAP1*, while in TN3 these mutations included *TP53*, *PLEKHA6*, *STXBP2* and *MED24*. In TN5, significant mutations were identified in *TP53*, *RGS7* and *NDST4* among other genes (Fig 4C). The CADD analysis showed that homozygous mutations in *TP53* had the most significant functional impact scores in all three tumors (Fig 4C). While most mutations in the three monoclonal tumors were detected in all of the tumor cells, there were a few mutations identified that occurred at lower frequencies, including *MED24*(4.8%) in TN3 and *OTUD5*(3.2%) and *PRDM5*(2%) in TN5; as well as *SENP6*, *MARCH6*, *PODXL2* and *ADCY8* (all at 1.8%) of cells in TN5, suggesting that these were later lineage events that emerged during tumor progression.

**Figure 4.**
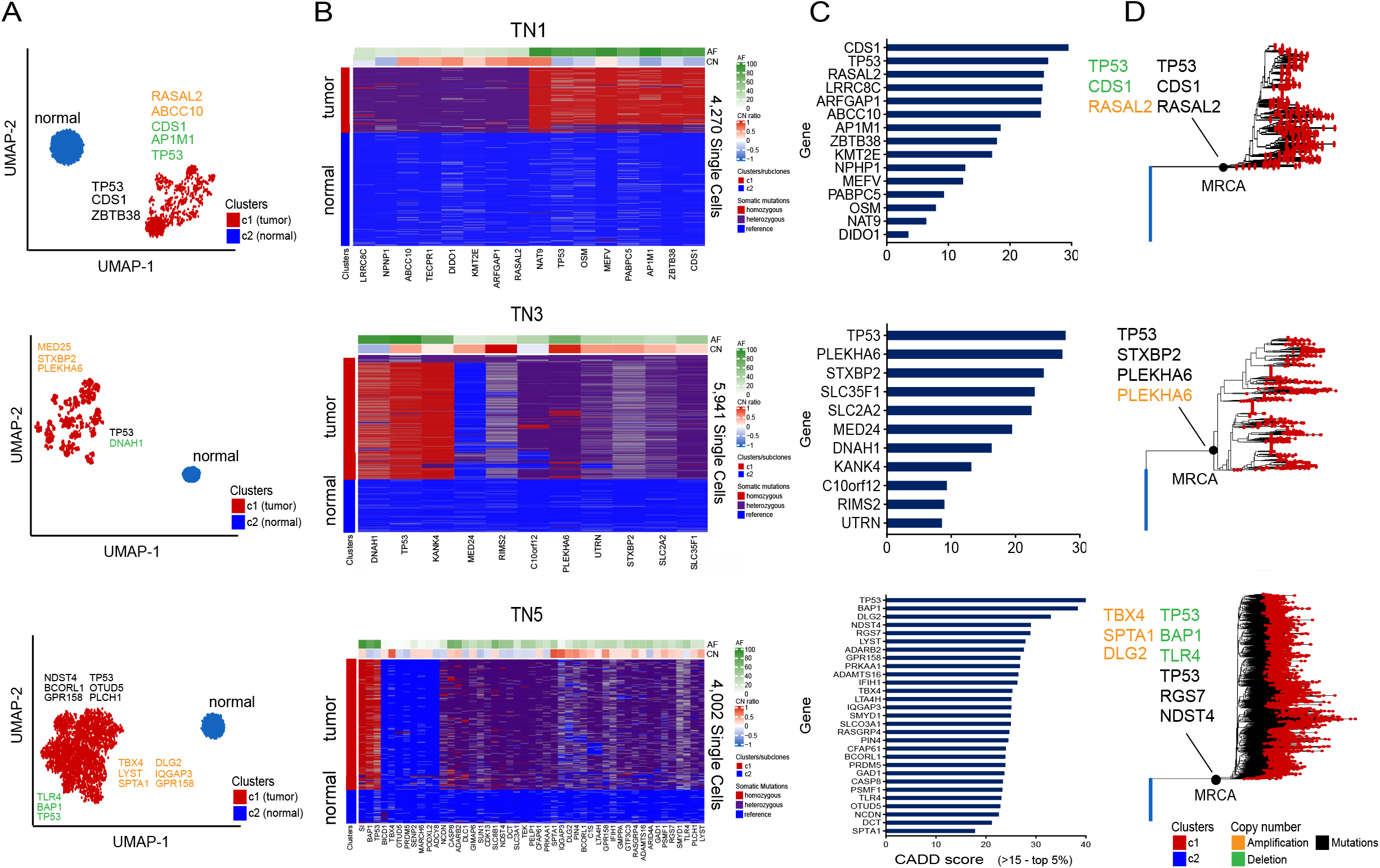
Clonal Diversity and Mutational Lineages of Three Monoclonal Tumors. (A) High-dimensional clustering using UMAP of 4,270 (TN1), 5,941 (TN3) and 4,002 (TN5) single cells, respectively, identified one major tumor cluster and one cluster of normal cells in each sample (B) Hierarchical clustering heatmaps of the mutations in TN1, TN3 and TN5 showing a monoclonal population of tumor cells, with cluster-level copy number estimations for each mutation shown in the header bar (C) CADD score is used to rank mutations’ predicted deleteriousness impact (CADD > 15, top 5%) in TN1, TN3 and TN5 (D) Phylogenetic reconstruction of the TN1, TN3 and TN5 mutation lineages, chronology, and timing of copy number aberrations relative to MRCA using a neighbor-joining tree with the MRCA annotated.

Targeted copy number inference showed that homozygous mutations *TP53*, *CDS1*, *AP1M1*, *ZBTB38*, *PABPC5* and *OSM* were deleted in TN1, while *PLEKHA6* was amplified in TN3 and *TBX4*, *DLG2* and *SPTA1* were amplified in TN5 (Fig. 4B). Neighbor-joining (NJ) trees were constructed to infer the chronology and order of SNVs and CNAs which showed that most events were truncal and occurred prior to the MRCA, after which the tumor cells expanded to form the tumor mass in each patient (Fig. 4D).

### Comparison of clonal substructure from single cell and bulk data

To understand the advantages of the MPT method for resolving clonal substructure, we directly compared bulk DNA-seq exome data to the single cell data in 5 patients. To asses the accuracy of the copy number data, we first calculated a consensus of the single cell data and compared it to the bulk exome data, which we considered to be the ‘gold standard’ reference for each patient (Fig 5A). Calculation of the Pearson correlation coefficients showed high correlation values (mean = 0.871) across the patients, suggesting that the single cell read count data, when merged together across cells accurately reflected the bulk copy number profiles. Notably, all of the high-level amplifications were detected in the single cell data from TN1 (*RASAL2*, *NAT9*), TN2 (*USP21*), TN3 (*RIMS2*), TN4 (*GRIN3A*), and TN5 (*IQGAP3*, *SPTA1*, *TBX4*, *PIN4*). Next, we compared the variant allele frequencies (VAFs) of the bulk exome data versus the combined single cell mutation frequencies (Fig 5B). Calculation of the Pearson correlation coefficients of the VAFs between the two datasets showed a high correlation (mean = 0.876) across the 5 patients, suggesting that they were highly concordant.

**Figure 5.**
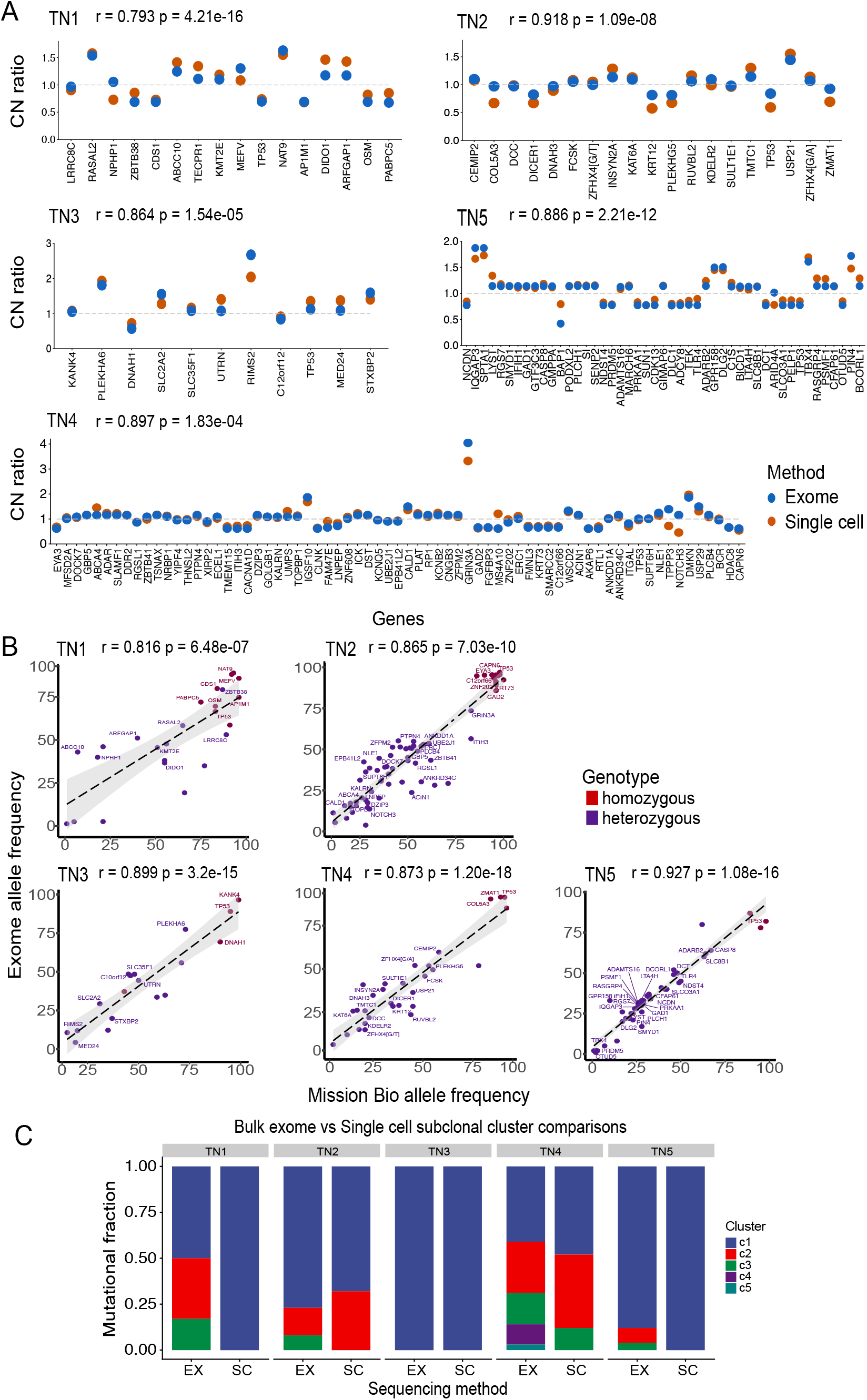
Comparison of MPT Single Cell vs Bulk Exome Sequencing Data. (A) Copy number states for each gene were estimated across all tumor cells from each patient using the Mission Bio data and compared to a pseudo-bulk single cell consensus copy number reference from WGS data (Pearson correlation) (B) Average allele frequencies for each mutation calculated across all single cells from each patient in the MPT data are compared to the bulk exome reference (Pearson correlation) (C) Pyclone2 subclone clustering of mutation frequencies from bulk exome data compared to subclone clusters detected from MPT scDNA-seq profiling in each patient.

Finally, we investigated whether the clonal composition of the bulk exome data could be estimated by clustering VAF distributions using PyClone2 (Roth et al. 2014) (Fig 5C). This data showed that PyClone2 often overestimated the number of subclones present in each tumor, in which the two monoclonal TNBC tumors (TN1, TN2) were estimated to have 3 subclones by PyClone2 and one monoclonal tumor (TN4) was shown to have 1 subclone. For the two polyclonal tumors, Pyclone2 estimated a higher number of subclones in the exome data of TN3 (3 by single cell, 5 by exome) and in TN5 (2 by single cell and 3 by exome). These data highlight the challenge in estimating the number of subclones from bulk-exome data based on VAFs, which single cell mutation data can accurately resolve.

## Discussion

Here, we report a new approach (MPT) that involves first performing deep-exome sequencing across a cohort of patients (eg. 5-10) and identifying mutations that can be pooled together to develop a custom capture platform that targets only these genomic regions for sequencing in scDNA-seq experiments. Our study utilized a microdroplet-based scDNA-seq platform (Mission Bio) to perform high-throughput analysis of thousand cells in parallel from 5 TNBC patients. By targeting regions with somatic mutations (identified *a priori* by bulk exome sequencing), we were able to profile 66 mutations in thousands of single cells using MPT. In contrast, previous studies that have used the same microdroplet platform (Mission Bio) to profile AML patients with pre-defined tumor panels, typically only measure a very small set of mutations (~1-8) in each patient across many cells (Pellegrino et al. 2018). From this data, we obtained sufficient genomic markers in each patient to resolve clonal substructure, reconstruct clonal lineages and identify driver mutations during breast tumor progression.

Our study showed that 3 tumors were mono-clonal (TN1, TN3, TN5), while 2 tumors (TN2, TN4) were multi-clonal out of 5 patients analyzed. We detected *TP53* mutations in all 5 TNBC patients, which we inferred to be one of the earliest truncal mutations that was acquired during tumor evolution. Additionally, our FACS data shows that genome doubling occurred in all 5 TNBC patients as evidenced by increased DNA ploidy (>2.65N), which is consistent with other studies in TNBC (Minussi et al. 2021). However, in addition to *TP53*, several additional early mutations and CNAs were also detected in the two polyclonal tumors. In TN4, early truncal homozygous mutations (*GRIN3A*, *PTPN4*, *AKAP6*) along with copy number losses (*TP53*, *NOTCH3*) and gains (*DNMK*, *GRIN3A*) co-occurred with *TP53* in the MRCA at the earliest stages of tumor progression. In TN2, early truncal homozygous mutations (*ZMAT1*, *SULTIE1*) and copy number losses (*TP53*, *ZMAT1*, *COL5A3*) and gains (*USP21*, *TMTC1*, *ZFHX4*[G/A]) co-occured with *TP53* mutations in the MRCA at the earliest stage of progression. These data are consistent with a punctuated copy number evolution (PCNE) model, in which short bursts of genome instability give rise to multiple clones that stably expand to form the tumor mass as previously reported in TNBC (Gao et al. 2016; Minussi et al. 2021). However, our data further suggest the possibility that mutations may also co-occur in short bursts of evolution, followed by stable clonal expansions that form the tumor mass in TNBC patients.

Our findings on very limited numbers of gradual intermediate mutations occurring between the lineages of subclones in the polyclonal tumors, and that there are few mutations occurring outside of the dominant clones in the monoclonal tumors is very unexpected. This data suggests either: 1) that the mutations were acquired simultaneously in short bursts of evolution, and that the mutation rate is very low outside of these events, or 2) that there are large selective sweeps leading to dominant clones that outcompete the other clones that harbored the gradual intermediate mutations. From our current data that is derived from a single time point, we cannot accurately distinguish between these two scenarios. Another notable limitation is that the MPT approach focused on profiling only targeted mutation sites in exons that were identified by bulk exome sequencing and did not perform unbiased profiling of mutation sites across the whole genome, possibly missing gradual intermediate mutations that may have occurred in single cells in non-coding and intergenic regions.

To better understand the concordance of MPT to bulk exome sequencing or WGS, we directly compared these independent datasets from the same tumors in 5 patients. Comparison of VAFs between the bulk and MPT merged single cell data showed a very high concordance (0.876, avg. Pearson correlation), confirming the accuracy of the MPT. Copy number estimation was performed on consensus clusters of the single cell MPT data for each mutation and compared to a pseudo-bulk derived reference. Even though the MPT estimation was limited due to coverage of only one amplicon per SNV, the copy number estimations showed a high concordance as well (avg. Pearson correlation – 0.871) when performed on groups of cells from each subclone cluster. This data allowed us to obtain both mutation information and copy number states for each subclone in the TNBC tumors, for reconstructing their order in the clonal lineages during tumor evolution. Our data also suggests that PyClone2 generally overclustered the bulk exome data and estimated higher number of subclones in each tumor compared to the scDNA-seq experiments. This analysis highlights the important of using single cell sequencing approaches to accurately resolve clonal substructure in solid tumors.

This study has several notable limitations. First, since this project was primarily for the development of the MPT approach, only 5 TNBC patients were analyzed. Thus, our biological conclusions regarding the evolutionary history of the tumors may not be generalizable across all TNBC cancers. Additionally, the custom amplicon panel included a limited number of targeted mutation sites (N=330), however future developments of the microdroplet technology (Mission Bio) may expand the number of amplicons to thousands of sites in the future. Another limitation was that the copy number measurements were not based on whole-genome measurements but were instead restricted to the targeted genomic regions covered by the mutations, thus limiting the overall genomic resolution of detection.

Important future directions will include applying MPT to study early-stage breast cancers such as atypical ductal hyperplasia (ADH) or DCIS to better understand mutation bursts and early events that initiate breast cancers in larger cohorts of patients. Additionally, it will be important to link clinical data on drug responses, survival and progression to the amount of intratumor heterogeneity in invasive breast cancers and other cancer types. In closing, the MPT approach provides a powerful new tool to target only information-rich genomic regions for scDNA-seq analysis to study clonal substructure and lineages across diverse solid cancer types.

## Data Access

The data has been deposited to the NCBI Sequencing Read Archive (SRA) under the accession: PRJNA763862

## Author Contributions

J.L. performed experiments, data analysis and prepared the manuscript. E.S. performed FACS and exome experiments. M.H. performed data analysis. F.M.B. obtained clinical tissues and clinical data. N.N. managed the project and wrote the manuscript.

## Acknowledgements

This study was supported by grants to N.E.N. from the American Cancer Society (129098-RSG-16-092-01-TBG), the NIH National Cancer Institute (RO1CA240526, RO1CA236864), the Emerson Collective Cancer Research Fund and the CPRIT Single Cell Genomics Center (RP180684). N.E.N. is an AAAS Wachtel Scholar, Andrew Sabin Family Fellow, Jack & Beverly Randall Innovator and AAAS Fellow. This study was supported by the MD Anderson Sequencing Core Facility Grant (CA016672), and MD Anderson T32 Translational Genomics and Precision Medicine Fellowship (CA217789). We thank Hongli Tang, Louis Ramagli and Erika Thompson for their help with next-generation sequencing. We are grateful to Alexander Davis for providing guidance on the mutation trees and Darlan Minussi, Naveen Ramesh, Tapsi Kumar, and Aislyn Schlack for data analysis support. Finally, we thank Robert Durruthy, Kelly Kaihara and Anjali Pradhan from Mission Bio for their computational and technical support for this project

## Methods

### Experimental Subject Details

We selected five snap-frozen tissue samples from women with untreated intraductal carcinoma breast cancer, including paired normal adjacent breast tissues, that were collected from the University of Texas MD Anderson Cancer Center Breast Tissue Bank. The patients were chosen based on their tumor grade (stage 3), age (37-71), histopathology, lack of treatment, tumor ploidy, and triple-negative (ER, PR, Her2) receptor status (Supplementary Table 1). IHC confirmed the ER and PR status of each tumor to be <1%, while Her2 negative status was determined by IHC or FISH cytogenetic analysis. All of the tissues were collected under informed consent, and the IRB for this study was approved by the MD Anderson Cancer Center Review Board.

### Generation of Nuclear Suspensions from Frozen Tissues

For each bulk (tumor and normal) and single cell experiment, nuclear suspensions were generated from frozen tissue using an NST/DAPI buffer (800 mL of NST [146 mM NaCl, 10 mM Tris base at pH 7.8, 1 mM CaCl2, 0.05% BSA, 0.2% Nonidet P-40, and 21 mM MgCl2]), 200 mL of 106 mM MgCl2 and 10 mg DAPI (Wang et al. 2014). To prepare the nuclear suspensions, sections of the tumors were minced with a surgical blade in petri dishes containing NST/DAPI, and then nuclei were filtered through a 40 um mesh into 1.5 mL Eppendorf tubes. Cell counts were obtained from DAPI utilizing the Countess (Invitrogen).

### FACS Enrichment of Aneuploid Nuclei

In order to enrich tumor nuclei from the tissue sections, millions of DAPI-stained nuclei were flow-sorted on the FACS Aria II (BD Biosciences). Nuclei were selected from the aneuploid distributions based on their DAPI intensity (>2N) relative to the diploid peaks of nuclei (2N and deposited in 1.5 ML Eppendorf tubes. The enriched aneuploid tumor nuclei suspensions were used for downstream single cell DNA-seq and for bulk exome profiling.

### Bulk Exome Sequencing

After using FACS to enriched aneuploid nuclear suspensions or matched normal tissues, the nuclei were prepared for sequencing by first fragmenting the DNA to 250 bp using a Covaris Sonicater and subsequently purifying with the Zymo DNA Clean & Concentrator Column kit (Zymo). Next, the fragmented DNA was barcoded using the NEBNext end repair model (NEB), dA-tailing (NEB), and quick ligation (NEB); and then library amplification was performed by PCR using NEBNext HiFi2x PCRmix. An exome capture was then applied using the Nimblegen SeqCap EZ Exome V2 kit (Roche), and both normal/tumor matched samples were sequenced paired-end at 100 bp on the Illumina HiSeq4000.

### Detection of Mutations in Bulk DNA Samples

The resulting FASTQ files (Supplementary Table 2) from the Hiseq4000 sequencing run of normal/tumor matched samples were then demultiplexed using our custom software code (deplexer.pl). Bowtie 2 (Langmead and Salzberg 2012) was then applied to align each FASTQ to the human genome reference (hg19) and then converted to individual BAM files by SAMtools (Li et al. 2009). Picard was used to remove PCR duplicates from the resulting BAM files. Indel regions were then realigned using the Genome Analysis Toolkit (GATK) (McKenna et al. 2010), and sequencing reads were filtered at a standard mapping quality score of 40. BEDTools (Quinlan and Hall 2010) was utilized to calculate coverage depth and breadth. The full in-house pipeline and scripts outlining each step can be downloaded from *Nature Protocols* (Leung et al. 2016). GATK was then applied with default parameters to detect variants and recalibrate quality scores resulting in a final bulk VCF file summarizing the results. Mutations were filtered out based on two distinct criteria: consensus filtering (mutations must be detected in at least 3 cells) and clustered regions (multiple mutations detected in a 10-bp sliding window). The resulting variants were then annotated by applying ANNOVAR to the VCF files using default parameters (Wang et al. 2010). Sites having low coverage (<10X) were annotated as missing values (NA) and nonvariant sites were labeled as germline reference.

### Mission Bio Custom Panel

Mission Bio’s custom design targeted panels allow the user to choose single nucleotide variants to be profiled over specific amplicon regions for high-throughput single cell DNA sequencing. Amplicon-based targeted sequencing is then applied to profile SNVs and estimate copy number variation (CNV) variation in up to 10,000 cells. In order to capture the most variance and heterogeneity in our tumor samples, we selected variants with allele frequencies ranging from 0.1% to 100% to capture the heterogenous subclonal events. Of the 379 genes submitted to Mission Bio for our custom design targeted panel, a final of 330 genes were included in the final panel construction. The final custom panel was then used on the Mission Bio Tapestri platform to profile SNVs utilizing high-throughput DNA sequencing.

### Mission Bio Single Cell DNA Sequencing

The MPT sequencing method utilizes the Mission Bio Tapestri micro-droplet platform to perform targeted high-throughput single cell DNA sequencing, where we followed the basic Mission Bio Tapestri default guidelines (Mission Bio User Guide). After isolating single cells in NST/DAPI buffer from the frozen tissue, we input 2-4,000 cells/uL into the Mission Bio Tapestri cartridge, where single cells are individually partitioned into nano-droplets with lysis buffer (Mission Bio) and incubated at 50C for 60 min on the PCR block (Bio Rad) to release the DNA. The DNA is then re-loaded into the Mission Bio cartridge, where barcoding beads and PCR reagents are combined in a second merged encapsulation. UV light (Analytik Jena XX-15L) is applied for 8 min to release the barcoded DNA from the beads, and the DNA is amplified via multiplexed PCR (Bio Rad) within the droplets (Mission Bio). The droplet emulsions are then broken and the DNA is extracted and purified using Ampure XP Beads (.72X). Qubit Fluorescence Quantification (Invitrogen) was used the quantify the concentration of DNA at an expected range of (0.2-4 ng/uL). For library construction, the i5 and i7 indexes are added and library amplification is performed (Mission Bio) on the PCR block (Bio Rad). The library is then purified using Ampure XP beads (0.69X) for an expected on-target size range of (350-550 bp) at a range of concentration (2-20 ng/uL). Qubit is used as a preliminary raw quantification of DNA concentration followed by the TapeStation (Agilent) for a more detailed view of the fragment size distribution in addition to the targeted concentration. The library was then diluted to 5nM (0.9-1.3 ng/uL) and sequenced on the Illumina HiSeq4000 at 150 paired-end.

### Mission Bio Mutation Detection Pipeline

The resulting single cell demultiplexed FASTQ files (Supplementary Table 3) were input into the Tapestri Sequencing Pipeline (Mission Bio). Adapter sequences are first trimmed from the raw sequencing reads using Cutadapt (Martin 2011, Bolger, Lohse et al. 2014). Short reads less than 30 nt are discarded and barcodes are extracted from the reads. Error correction is used (Hamming distance or Levenshtein distance) to correct barcodes with a partial match to increase yields. A cell calling algorithm is then applied to only select cells that have at least 80% amplicon read completeness and pass a total reads cutoff. The reads are then mapped to the reference genome (hg19) using the BWA-MEM algorithm (Langmead, Trapnell et al. 2009; Kim, Pertea et al. 2013) with default parameters discarding any unmapped reads. The cells are genotyped using the GATK with a joint calling approach that follows GATK Best Practices (DePristo, Banks et al. 2011; Van der Auwera, Carneiro et al, 2013). Finally, joint genotyping is performed for all cells using GATK’s GenotypeGVCFs tool, VCF and HDF5 files are generated, and the genotypes and cell matrix are converted into an open-source loom format for further analysis (Zeisel, Hochgerner, et al. 2018). Loom files were input into Mission Bio Tapestri Insights, where default parameters were used to filter the data by cell quality (<30), read depth (<10), alternate allele frequency (<20%), variants genotyped/cell (<50%), percent genotypes present (<50%) and percent cells mutated (<1%). Heterozygous germline mutations present in more than 95% of cells were also removed, and amplicons and/or cells that had NA values in over 50% of cells were excluded.

### Clustering and Doublet Removal

PCA (Pearson 1901) was employed to reduce the high dimensional space, and UMAP (McInnes et al. 2018) was further applied to the PCA selected features to partition the cells in distinct clusters where outliers (eg. low quality, depth, NAs) were further excluded. Unsupervised hierarchical clustering via ComplexHeatMap (Gu et al. 2016) was used define and visualize the final subclones based on the determined features (linkage=complete, distance= Euclidean), which were then overlaid on the UMAP projection for annotation. Single cell doublets clusters were identified for removal by analyzing coverage depth comparisons against all remaining subclone cluster mutation depths. Specifically, median coverage read depth counts were calculated across all mutations for each cell per cluster, then normalized across all amplicons by the normal diploid population read counts to assign an effective ratio value for each cluster. The (log2) absolute values of the normalized read counts (effective ratio values) for each cluster were compared and each significantly higher doublet cluster was removed (Supplementary Figure 2).

### Estimation of Mutation Impact

To predict the functional impact of the detected variants, Combined Annotation-Dependent Depletion (CADD) (Kircher et al. 2014) was coupled with POLYPHEN2 (Adzhubei et al. 2010) and SIFT (Vaser et al. 2016) to prioritize causal variants based on over 60 combined genomic features. Most annotations tend to exploit a single information type and/or are restricted in scope, so a broadly applicable metric that objectively weights and integrates diverse information was needed. Thus, CADD integrates multiple annotations into one metric by contrasting variants that survived natural selection with simulated mutations. For each sample, the mutations were first filtered by a POLYPHEN or (1-SIFT) score over 0.8, then the top 30 deleterious genes were selected from this filtered pool based on the CADD functional impact scores for each patient scaled by CADD > 10 = top 10%, CADD > 20 = top 1%, and CADD > 30 = top 0.1%.

### Integrated Phylogenic Trees

Neighbor Joining (NJ) trees were constructed from the clustered SNV data using Ape (Paradis et al. 2004), and the cluster annotations defined through UMAP and hierarchical clustering were overlaid onto the resulting NJ trees. The respective mutations for each sample were annotated on the NJ tree based on the clusters in which they were identified. Estimated copy number aberrations for deleterious genes showing significant CNV per cluster were annotated on the trees based on the clusters in which they were identified.

### Copy Number Estimation from Mission Bio Data

Mission Bio Copy Number Analysis is a tool for estimating copy number from single cell read depth and was provided by Mission Bio. After clustering by mutations and normalizing for read depth across all cells and amplicons, the read depth for each cell per mutation is then divided by the median reference read depth for the same mutation, thereby providing an effective copy number ratio value for each cell per gene per cluster. This effective ratio value then serves as the estimated copy number. While this provides single cell copy number estimations per cell, the data has significant technical noise at single cell resolution. To correct for this, we calculated the median copy number for each gene per cluster, which was used to annotate hierarchal clustering heat maps, integrate NJ trees, and compare panel accuracy against bulk exome copy number data.

### Single Cell vs Bulk Exome Comparisons

The median copy number ratio values estimated from the Mission Bio data per gene were compared against the median bulk copy number ratio values. Correlation was calculated using Pearson. Next, the allele frequencies of each variant in the Mission Bio and bulk data were compared, and the relationship was defined by Pearson’s correlation. Pyclone2 (Roth et al. 2014) was applied to the bulk exome data using default parameters to estimate the total number and size of the clusters. The resulting Pyclone2 cluster assignments were then compared to the clusters defined by the single cell analysis (Supplementary Figure 1).

